# Pan-cistrome analysis of the leaf accessible chromatin regions of 214 maize inbred lines

**DOI:** 10.1101/2024.10.14.618191

**Authors:** Yongli Zhu, Heiyuen Ngan, Tao Zhu, Lili Nan, Wenqiang Li, Yingjie Xiao, Lin Zhuo, Dijun Chen, Xiaoyu Tu, Kang Gao, Jianbing Yan, Silin Zhong, Ning Yang

**Author notes:** Co-first authors (Yongli Zhu, Heiyuen Ngan). Correspondence: Silin Zhong; Ning Yang.

## Abstract

Characterizing the noncoding regions of the genome, particularly the cis-regulatory elements (CREs) located in accessible chromatin regions (ACRs) in gene promoters, is crucial for understanding how gene expression is regulated and how genotype contributes to phenotypic diversity in plants.In this study, we used ATAC-seq to resequence and map the ACRs of 214 maize inbred lines. We identified 82,174 ACRs and reported that 39.55% of them exhibited significant variation across the population. Next, we used the accessibility of ACRs as a quantitative feature and performed a chromatin associability GWAS (caGWAS), resulting in 27,004 caQTLs. Among them, 2,463 were predicted to disrupt the cis-regulatory elements that TF binds in these ACRs, suggesting that TF binding affects chromatin accessibility. Specifically, we identified two caACRs that regulate the expression of *fad7*, which encodes a fatty acid desaturase and affects the linolenic acid content.Our findings highlight the dynamic nature of maize ACRs and demonstrate that chromatin accessibility information could be used for GWASs, offering key insights into the genetic and regulatory mechanisms of chromatin accessibility and its impact on complex traits in plants.

## INTRODUCTION

For many years, the study of functional genomics in both animals (Mali et al., 2013; Wang et al., 2014) and plants (Huang et al., 2012; Tian et al., 2011) has focused primarily on the coding regions of the genome. However, the transcriptional regulatory mechanisms associated with noncoding regions and their contributions to trait variation remain underexplored. Recent advancements in the field of epigenomics (Satterlee et al., 2019) and the rapid development of next-generation sequencing technologies (Goodwin et al., 2016) have shifted the focus toward understanding the role of active regulatory regions within the noncoding portions of the genome. Techniques such as DNase-seq (Boyle et al., 2008; Hesselberth et al., 2009), MNase-seq (Schones et al., 2008), and ATAC-seq (Buenrostro et al., 2013) have enabled researchers to systematically identify the regulatory regions in gene promoters, specifically the accessible chromatin regions (ACRs) where transcription factors (TFs) bind. ATAC-seq, in particular, has garnered widespread adoption because of its procedural simplicity and temporal efficiency (Buenrostro et al., 2013). By employing these advanced techniques, scientists are beginning to dissect the complex mechanisms of gene regulation and phenotypic diversity, which can be explained by noncoding sequences.

The temporal and spatial pattern of gene expression is encoded within the genomic DNA. This information is read by TFs, which recognize and bind specific short DNA sequences known as cis-regulatory elements (CREs) within the ACRs located in gene promoters and distal enhancers. As part of the Human Encyclopedia of DNA Elements (ENCODE) project (Maher 2012), Thurman et al. generated the first chromatin accessibility map for the human genome, providing a valuable resource for studying the genetic architecture underlying diseases and other complex traits (Thurman et al., 2012). Unlike animal genes, where distal enhancers can be located hundreds of kilobases upstream or downstream of the transcriptional start site, enhancers in plant genomes are often found in closer proximity to the gene of interest, typically within the 5’ upstream region. For example, Hendelman et al. employed CRISPR/Cas9 to edit different regions of the 5’ upstream sequence of the tomato WOX9 gene randomly. Although the precise locations of the ACRs within this noncoding sequence are not known, their approach generated a range of distinct WOX9 expression patterns, leading to varying phenotypes in both embryo and floral development (Hendelman et al 2021). Similar strategies have yielded diverse phenotypic outcomes in other species, such as maize (Liu et al., 2021) and rice (Song et al., 2022; Zhou et al., 2023). Despite the success of random editing approaches, the ability to identify precise genomic regions or even specific DNA motifs as targets will soon emerge as a main challenge for improving traits through the use of targeted genome editing.

Over the past 20 years, high-density SNP genotyping and DNA resequencing have revealed the majority of genetic variation in many organisms. Genome-wide association studies (GWASs) have proven to be powerful tools for linking these variations to their corresponding genetic traits (Liu et al., 2019). In diverse maize populations, numerous associations have been established between genetic variation and a wide range of traits, including agronomic (Yang et al., 2014), transcriptional (Fu et al., 2013), and metabolic (Wen et al., 2018) characteristics. A population-scale study provides valuable insights into genetic variations affecting chromatin accessibility. Currint et al. demonstrated that genetic variations can disrupt the binding sites of transcription factors, resulting in either increased or decreased chromatin accessibility (Currin et al., 2021). Additionally, Gate et al. reported that 15% of genetic variations influence chromatin accessibility (Gate et al., 2018). These findings suggest that chromatin accessibility is modulated by genetic variations and can be regulated through alterations in transcription factor-binding sites.

Maize is an important crop with extensive genetic and genomic resources. Significant progress has recently been made in maize functional genomics, including the characterization of long-range chromatin interactions (Hi-C and ChIA-PET), DNA methylation, histone modification, single-cell gene expression, and chromatin accessibility. Additionally, large-scale TF binding data are now available for 104 TFs in leaf mesophyll cells (Tu et al., 2020), enabling systematic analysis of the CREs that regulate gene expression. However, most functional genomics studies have focused on the B73 cultivar, suggesting that variations in noncoding genes could account for nearly half of the additive genetic variance underlying phenotypic diversity (Rodgers-Melnick et al., 2016; Song et al., 2021). An understanding of how extant genetic variants affect cis-regulatory activity remains limited at the population scale. Therefore, investigating the diversity of chromatin accessibility across a diverse maize association mapping panel is crucial for elucidating how CREs contribute to the complex transcriptional regulatory mechanisms underlying phenotypic diversity.

In this study, we utilized ATAC-seq data from the leaf tissue of 214 maize inbred lines to conduct genome-wide association analyses. We also integrated ATAC-seq data with multiomic data, including ChIA-PET, ChIP-seq, RNA-seq, and phenotypic data, to systematically examine the role of chromatin accessibility in shaping the transcriptional landscape and complex traits. This study advances our understanding of the genetic basis of chromatin accessibility and contributes to unravelling the noncoding regulatory complexity associated with complex traits.

## RESULTS

### The accessible chromatin region varies widely across the maize association population

We performed ATAC-seq to profile chromatin accessibility in the seedling leaves of a diverse panel of 214 maize inbred lines. This panel was a subset of an association mapping panel (Yang et al., 2011) for which omics data had been generated (Gui et al., 2020). In total, we obtained approximately 9.46 billion reads, with an average of 43.99 million reads per sample. These reads were mapped to the B73 reference genome (RefGen_V4) (Jiao et al., 2017), with an average mapping rate of 83.56% (Additional file 2: Table S1). We used MACS2 (Zhang et al., 2008), a peak caller of the ENCODE ATAC-seq pipeline, to identify ACRs (see Methods). The number of ACRs identified per sample ranged from 5,554-77,030, with an average of 37,221 (Additional file 1: Fig. S1a). The FRiP (fraction of reads in ACRs) values ranged from 8.32--76.59%, with an average of 41.33%. In particular, 97.20% of the TSS enrichment values were greater than 4 (Additional file 2: Table S1). The above results indicate that high-quality libraries with high signal-to-noise ratios were generated from maize seedling leaves (Yan et al., 2020). A total of 36 samples had fewer ACRs (< 20,000), likely due to a lower amount of sequencing data or a reduced number of reads in the actual ACRs (Additional file 1: Fig. S1b). To ensure sample diversity and minimize bias from differences in sequencing quantity, low-frequency ACRs (< 15 in the association panel) were removed. We obtained 82,174 consensus ACRs, ∼44 Mb in total, representing 2.13% of the B73 genome. We quantified the consensus ACRs in the population by counting the number of cleavage sites within each ACR (see Methods). Principal component analyses (PCAs) revealed significant batch effects (Additional file 1: Fig. S1c). To address the batch effects, we utilized the COMBAT function from the SVA-R (Johnson et al., 2007) (see Methods). After correction, no discernible batch effect was observed (Additional file 1: Fig. S1d).

Promoters, enhancers and other cis-regulatory regions typically show enrichment of active histone modifications, such as H3K27ac, H3K4me1, and H3K4me3 (Buenrostro et al., 2013; Kouzarides 2007; Lara-Astiaso et al., 2014). Our 82,174 identified ACRs were located primarily at the ends of chromosomes, displaying a distribution pattern similar to that of genes and those with active histone modifications (Fig. 1a). Among these ACRs, 88.63% were shorter than 750 bp (Fig. 1b), and more than half were located within 10 kb of nearby genes (Fig. 1c). This finding is consistent with previous studies in other species (Maher et al., 2018), where the majority of ACRs are found upstream of the nearest gene (< 10 kb to the TSS). Additionally, an analysis of ChIP-seq data for 104 maize transcription factors in seedling leaf tissue (Tu et al., 2020) revealed that over 79.55% of the ACRs overlapped with transcription factor-binding sites (defined as covering 50% of the ACR length, Fig. 1d), suggesting that the identified ACRs are functional, as indicated by TF binding.

**Fig. 1.**
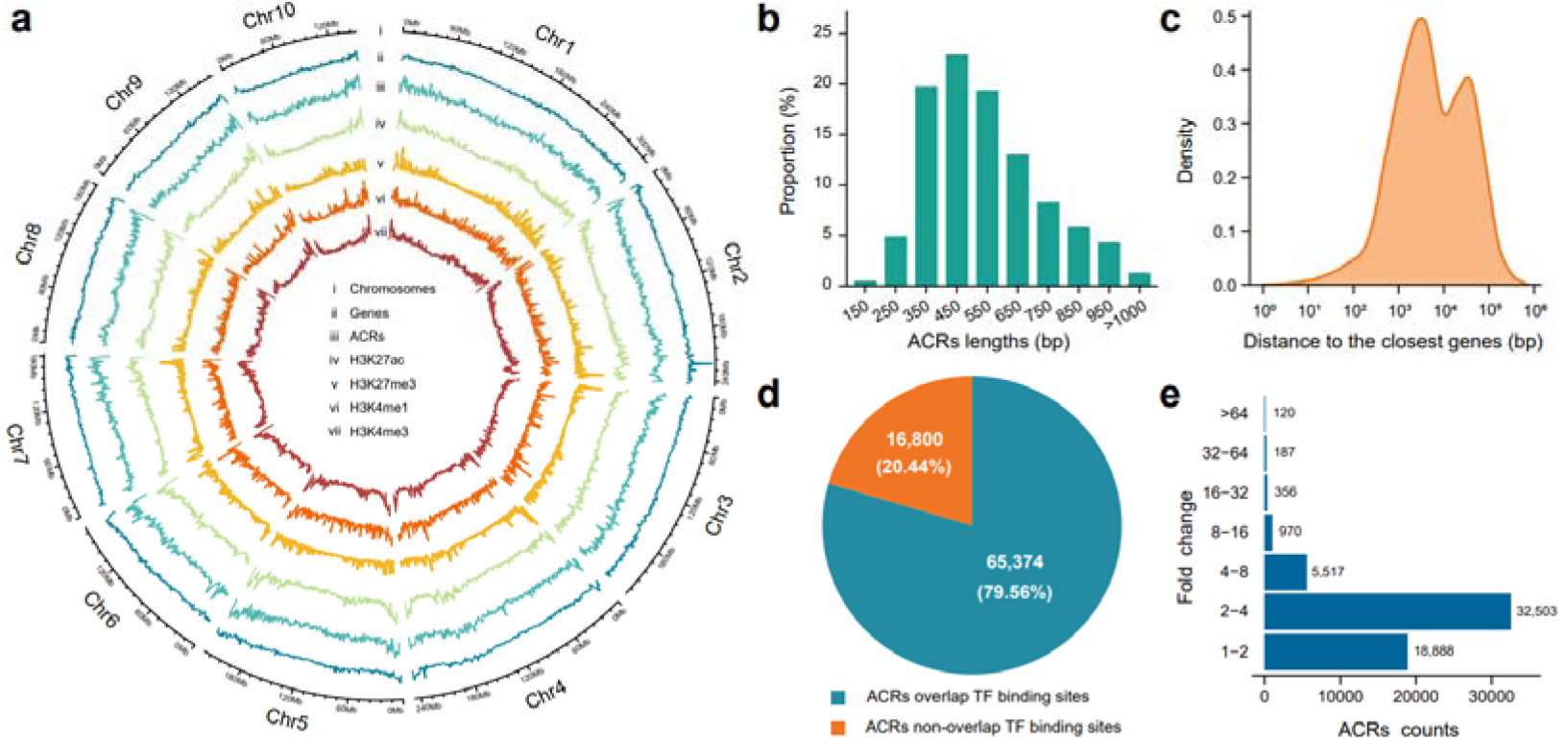
Basic characteristics of ACRs. **a** Distribution of genes, ACRs, H3K27ac, H3K27me3, H3K4me1, and H3K4me3 on chromosomes of B73. **b** Proportion of ACR length distributions. **c** Distance distribution between ACRs and the nearest gene. **d** Number of ACRs overlapping with transcription factor-binding sites. **e** Fold change in population chromatin accessibility variation of ACRs.

We also observed that chromatin accessibility varies considerably among maize inbred lines. A total of 16,541 ACRs were detected across all the inbred lines and were classified as shared ACRs (Additional file 1: Fig. S1e). Among these shared regions, 64.15% were annotated as proximal promoter ACRs (within 1 kb of the TSS, Additional file 1: Fig. S1f). In contrast, only 33.03% of the nonshared ACRs were located in the proximal promoter, suggesting that the ACRs in distal enhancers are more dynamic (Additional file 1: Fig. S1f). Across the population, 32,503 (39.55%) ACRs exhibited variability in chromatin accessibility ranging from 2 to 4 fold, with over 6,000 ACRs showing variation greater than 4-fold (Fig. 1e). These results suggest significant differences in chromatin accessibility at the population level.

### GWASs suggest that chromatin accessibility is regulated by genetic factors

We applied a standard genotyping pipeline (Pabinger et al., 2014) to ATAC-seq data from 214 samples and identified 349,492 single nucleotide polymorphisms (SNPs) (see Methods). Because ATAC-seq analysis was performed on leaf tissue, some genes with tissue-specific expression had no mapped reads in their promoter regions. This could lead to a loss of potentially informative SNPs compared with whole-genome sequencing data. Therefore, we also used the genotypes from a published dataset with over 10 million SNPs from 507 maize inbred lines as a reference panel (Chen et al., 2022) to impute missing genotypes. After imputation, we obtained 7,443,172 SNPs for 214 samples (see Methods).

We performed genome-wide association analysis (GWAS) via both ATAC-seq-only and imputed SNPs via a mixed linear model (see Methods). A total of 58,541 ACRs (present in 90% or more of 214 samples; Additional file 2: Table S2) were utilized for the association analysis. The GWAS identified 67,285 significant SNPs from the ATAC-seq-only dataset associated with 10,835 ACRs, whereas 1,894,877 significant SNPs from the imputed SNP dataset were associated with 18,428 ACRs. Among these, 9,392 ACRs were identified by both analyses (Additional file 1: Fig. S2a), accounting for 86.68% of the ACRs with ATAC-seq-only SNPs detected and 50.96% of the ACRs with imputed SNPs detected. These results suggest that imputing additional SNPs allows the detection of a greater number of genetically regulated ACRs.

Next, we focused on the GWAS results using imputed SNPs. Identifying 27,004 caQTLs (chromatin accessibility quantitative trait loci) for 18,428 ACRs (with caQTLs, termed caACRs). Among the caACRs, 15,522 caACRs (84.23%) had only 1 caQTL, 1,476 caACRs had 2 caQTLs, and 1430 caACRs had 3 or more caQTLs (Fig. 2a). caACRs were plotted against the peak SNPs, revealing strong enrichment along the diagonal, which indicates a strong local regulatory relationship with chromatin accessibility (Fig. 2b). SNPs near ACRs may affect chromatin accessibility by disrupting TF binding sites (Currin et al., 2021; Gate et al., 2018). Therefore, we focused on 5,948 (32.27%) caACRs and their proximal peak SNPs (within ±1 kb of the ACR, Fig. 2c). While GWAS approaches could identify individual loci with large effects, it is likely that genetic variations contribute smaller effect sizes to the degree of chromatin accessibility. We estimated that genetic variation accounted for approximately 36.43% of the additive genetic variance across the 58,541 ACRs analysed (see Methods). Notably, genetic variation contributed more to the additive genetic variance in caACRs (∼55.36%) than in noncaACRs (∼27.78%) (Fig. 2d).

**Fig. 2.**
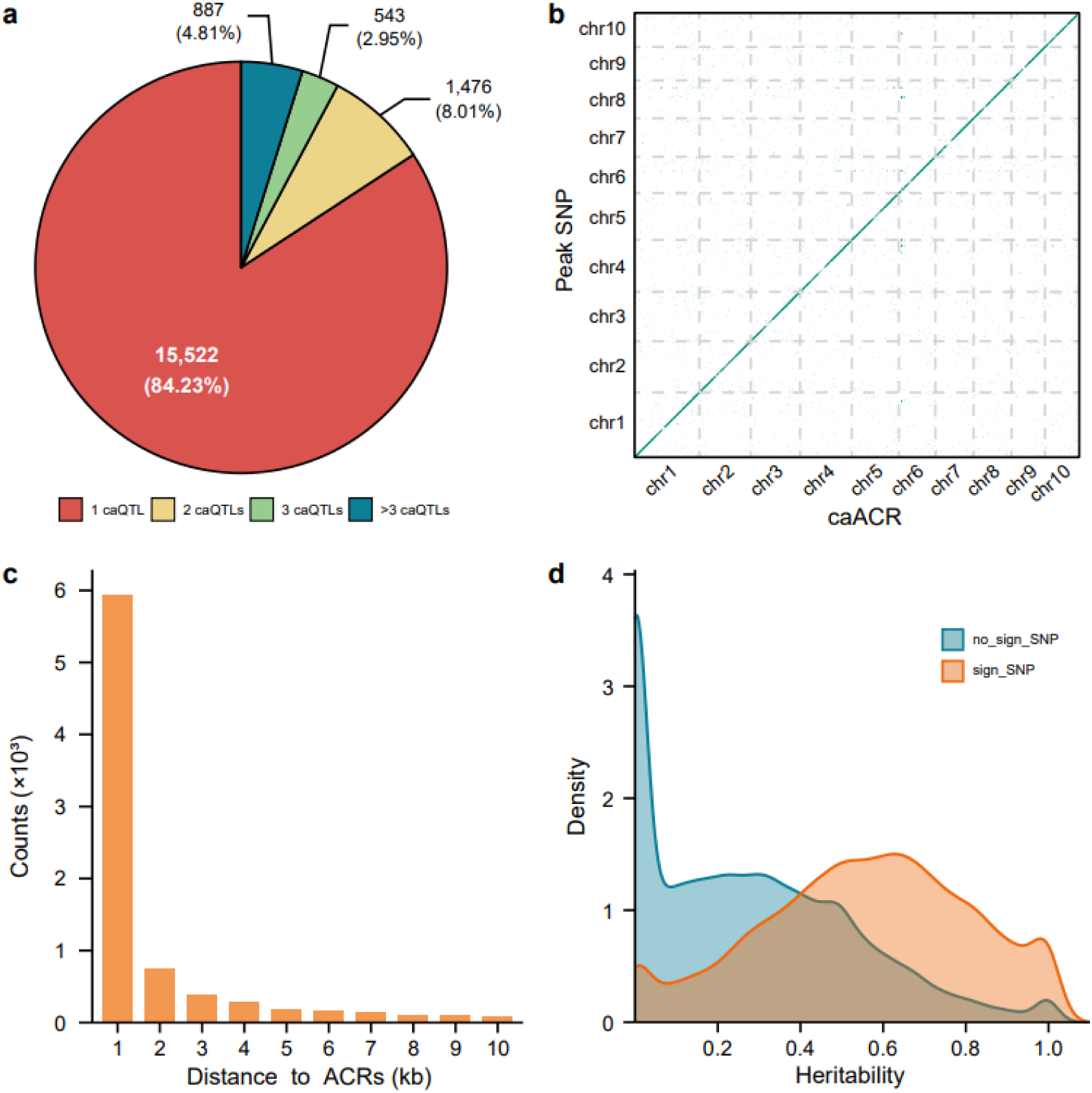
Genetic basis of ACRs. **a** The number of caQTLs associated with each ACR. **b** The start position of ACRs is plotted against the associated SNPs with the Bonferroni threshold. **c** Frequency distribution of distances between peak SNPs and ACRs. **d** Distribution of genetic heritability for ACRs with significant SNPs versus those without significant SNPs.

### Variation in chromatin accessibility associated with TF motif disruption

Human genetic variants located within caACRs have been shown to regulate chromatin accessibility by altering TF binding sites (Currin et al., 2021; Gate et al., 2018). Statistical analysis of the maize SNP genetic effects revealed that associated SNPs within caACRs tend to have greater effects than those outside (Fig. 3a). To test whether SNPs can disrupt TF binding sites, we selected 4,701 SNPs located within the ACRs that are the most significant SNPs. Additionally, we selected 870 associated SNPs located within ACRs that are in strong linkage disequilibrium (*r^2^* > 0.7) with the most significant SNPs. Using a total of 5,571 SNPs, we employed motifbreakR (Coetzee et al., 2015) to predict whether they could disrupt TF binding sites and affect chromatin accessibility (see Methods). To compare the proportion of peak SNPs within ACRs that disrupt TF binding motifs between ATAC-seq-only and imputed SNPs, we also predicted 3,261 peak SNPs within 2,929 ACRs identified via ATAC-seq-only SNPs.

**Fig. 3.**
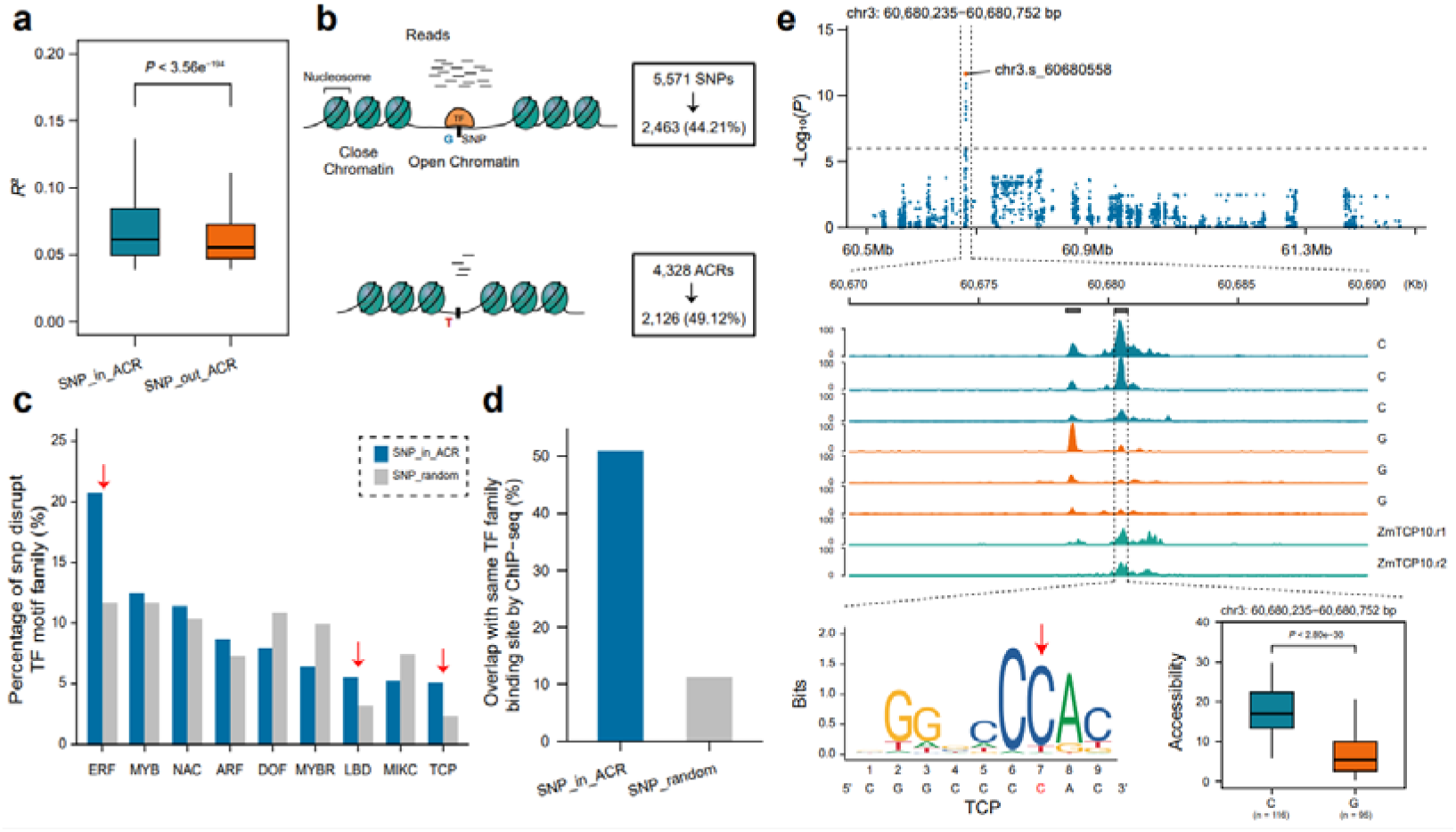
Mechanisms influencing chromatin accessibility. **a** Genetic effect of SNPs located within and outside ACRs. **b** Prediction of disrupted TF motifs (*P* < 1×10^-4^) by peak SNPs within ACRs or SNPs in strong linkage disequilibrium (*r^2^* > 0.7) with the peak SNP. **c** Enrichment analysis of SNP-disrupted TF families. **d** Proportion of SNP-disrupted TF families consistent with the same TF binding sites according to ChIP-seq. **e** Manhattan plot of chr 3:60,680,235--60,680,752 (top). The peak SNP (chr3.s_60,680,558) within the ACR is predicted to disrupt the TCP family motif with the red arrow (bottom left). The box plot displays the significant impact of the two genotypes on chromatin accessibility (bottom right). The genome browser track shows the difference in chromatin accessibility between the two genotypes and the ChIP-seq data, indicating that ZmTCP10 binds to the ACR region (center).

Among the 5,571 SNPs within 4,328 ACRs, 2,463 (44.21%, *P* < 1.0×10^-4^) SNPs within 2,126 ACRs (49.12%) altered the binding site of a TF motif (Fig. 3b). Among the 3,261 peak SNPs within 2,929 ACRs, 1425 (43.69%) SNPs within 1342 ACRs (45.81%) altered the binding site of a TF motif. There was no significant difference between ATAC-seq-only and imputed SNPs (43.69% vs 44.21%, *P* = 0.36, Additional file 1: Fig. S2b), indicating that the impact of SNPs detected from ATAC-seq data is equivalent to that of imputed SNPs. Among the 2,126 ACRs, 686 (32.26%) contained at least one SNP predicted to disrupt a motif, and 1,440 of these ACRs contained 2 or more predicted motif-disrupting SNPs. The disrupted motif families were significantly enriched in the ERF, LBD and TCP TF families (permutation FDR < 0.05, Fig. 3c).

To test whether the predicted TF binding-disrupting SNPs are real, we examined the published 104 TF ChIP-seq datasets and found that 1,252 (50.83%) of the predicted SNPs overlapped with ChIP-seq peaks of the same TF families, which was significantly greater than expected at random (permutation FDR < 0.05, Fig. 3d). These findings demonstrated that the predicted motif-disrupting SNPs are likely the binding sites of the corresponding TF families. For example, we observed a significant difference in chromatin accessibility (chr3:60,680,235-60,680,752) between the two alleles at a peak SNP (*P* < 2.80×10^-30^, N = C/G = 116/95, Fig. 3e). The peak SNP was predicted to disrupt a TCP binding motif (*P* < 1.90×10^-5^), and the ChIP-seq peak confirmed the presence of a binding site for ZmTCP10 in this region (Fig. 3e). Similarly, a peak SNP (chr10.s_139136345) was significantly associated with chromatin accessibility (chr10:139,138,162-139,138,791, *P* < 3.97×10^-50^, N = G/T = 156/59). This SNP was also predicted to disrupt a MYB binding motif (*P* < 1.52×10^-5^) and coincided with a binding peak for the TF ZmMYB81 (Additional file 1: Fig. S3).

### Many long-range regulatory elements may exist in the maize genome

To explore the biological functions of chromatin accessibility variations, we conducted an integrative analysis combining RNA-seq (Wu et al., 2021), ChIP-seq (Tu et al., 2020), and ChIA-PET (Peng et al., 2019) data. The challenge of connecting ACRs to their target genes was addressed via two different approaches (see Methods). First, we examined the colocalization between caQTLs of the caACRs and the eQTLs of genes. Second, we employed transcriptome-wide association studies (TWASs) to assess the relationship between chromatin accessibility and gene expression (Gusev et al., 2016). Among the 5,948 caACRs (defined as ±1 kb around peak SNPs), caQTLs of 452 caACRs (7.60%) were found to be in strong LD with at least one eQTL of eGenes (with eQTLs termed eGenes), resulting in the identification of 478 caACR-eGenes. Additionally, TWAS revealed that 1,508 caACRs were significantly associated (BH rejection threshold: *P* < 10^-6^) with 1,510 genes, yielding 1,934 caACR-gene pairs. Notably, 422 (88.28%) of the caACR gene pairs identified by colocalization were consistent with the results from TWAS (Fig. 4a).

**Fig. 4.**
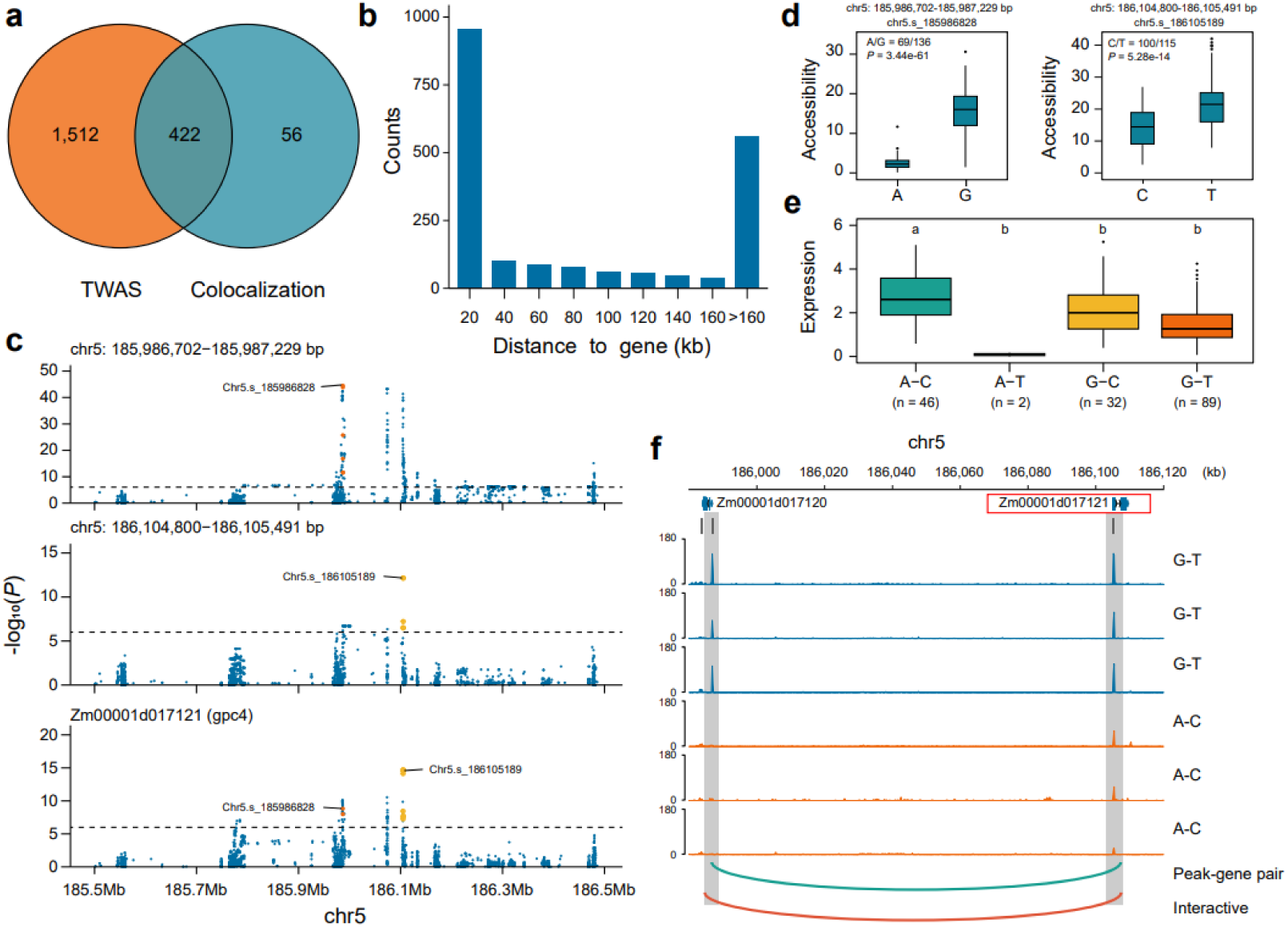
ACRs regulate gene expression. **a** Two approaches, TWAS and colocalization, for identifying ACR-regulating genes. **b** Distribution of distances between 1,990 pairs of caACRs and genes. **c** Local Manhattan plots of chr5:185,986,702-185,987,229, chr5:186,104,800-186,105,491, and *gpc4*. The red dots are significant SNPs within chr5:185,986,702--185,987,229, and the orange dots within chr5:186,104,800--186,105,491. The labels represent peak SNPs of ACRs. **d** Two genotypes of peak SNPs in two ACRs significantly impact chromatin accessibility. **e** Four genotypes of peak SNPs significantly impact *gpc4* expression. **f** Genome browser track displaying differences in the chromatin accessibility of the two ACRs in the two genotypes. The green line represents the association between ACRs and *gpc4*, and the black line represents the chromatin interaction between them.

In total, we identified 1,990 pairs of caACRs and genes, which consisted of 1,555 unique caACRs and 1,555 unique genes. Among these genes, 78.91% (1,227) of the caACRs regulated only one gene, whereas only a small fraction (80/1,990) regulated three or more genes (Additional file 2: Table S3). Among the 1,990 pairs, 933 (46.8%) involved caACRs located more than 40 kb from their associated genes (Fig. 4b), and 90 pairs were supported by chromatin interactions. These findings underscore the widespread presence of long-range regulatory elements in maize.

We also identified a distal caACR regulating the expression of *gpc4* (3-phosphoglycerate dehydrogenase) (Hou et al., 2022), a key enzyme in the glycolysis pathway. caQTLs located at chr5:185,986,702-185,987,229 and chr5:186,104,800-186,105,491 colocalized with eQTLs of *gpc4* (Fig. 4c). The peak SNPs (chr5.s_185986828 for chr5:185,986,702--185,987,229, chr5.s_186105189 for chr5:186,104,800--186,105,491) significantly influenced chromatin accessibility (Fig. 4d). The A-C haplotype presented the highest *gpc4* expression levels (Fig. 4e), suggesting a negative correlation between chromatin accessibility and gene expression. These two caACRs may act as repressive elements. Notably, *gpc4* is regulated by two caQTLs, one proximal (chr5:186,104,800--186,105,491, < 1 kb) and one distal (chr5:185,986,702--185,987,229, 117 kb). The distal caACR was found to interact with *gpc4* through chromatin interactions (Fig. 4f), suggesting that this long-range regulatory element can regulate *gpc4* expression.

### Chromatin accessibility is important for the regulation of complex traits

To investigate the potential associations between caACRs and agronomic and metabolic traits, we analysed QTLs from previously published studies via the same association mapping panel (Gui et al., 2020). In total, we identified 1,555 caACRs located in agronomic or metabolic trait QTLs. These caACRs were significantly enriched in QTLs related to agronomic traits, such as flowering and yield, as well as metabolic traits, including amino acids, sugars, and fatty acids (P < 0.05, Fig. 5a).

**Fig. 5.**
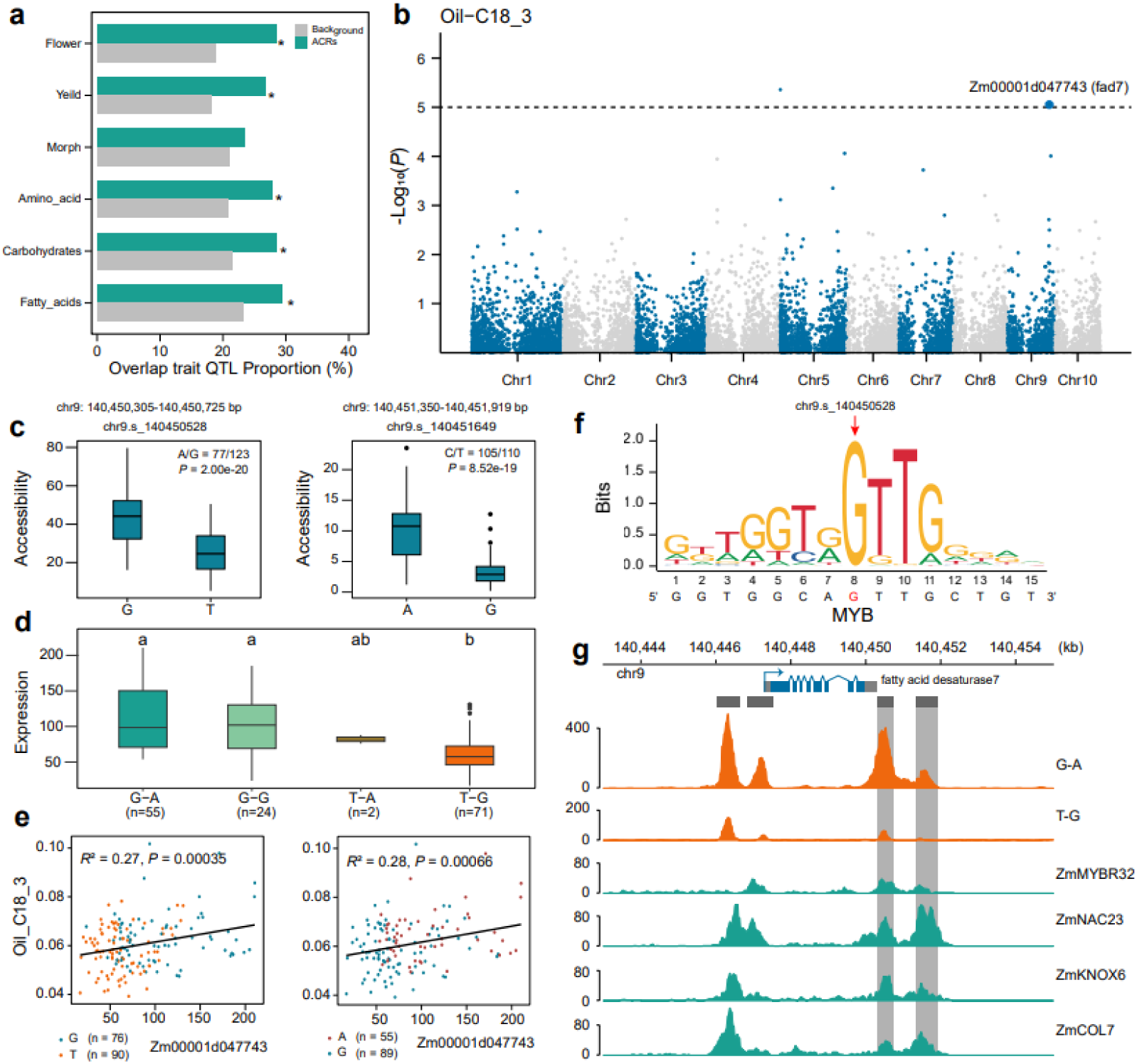
ACRs regulate complex traits. **a** Proportion of GWAS (including flowering, yield, and morphology, as well as amino acid, carbohydrate, and fatty acid metabolism) QLT for an assortment of traits overlapping with caACRs. Traits in which the enrichment was statistically significant (*P*_<_0.05) are labelled with an asterisk. **b** Manhattan plot of the TWAS between Oli-C18:3 and *fad7* expression. **c** G/T genotypes of chr9:140,450,305-140,450,725 and A/G genotypes of chr9:140,451,350-140,451,919 impact chromatin accessibility. **d** Four genotypes of peak SNPs significantly impact *fad7* expression. **e** G/T genotypes of chr9:140,450,305-140,450,725 and A/G genotypes of chr9:140,451,350-140,451,919 impact *fad7* expression and Oli-C18:3 content. **f** The peak SNP (chr9.s_140450528) within the ACR is predicted to disrupt the MYB family motif with the red arrow. **g** Genome browser track showing the difference in chromatin accessibility between the two genotypes (orange) and the binding of TFs to these ACRs (green).

Through TWAS analysis, we found a strong association between the expression of *fad7* (fatty acid desaturase) and the linolenic acid content (Li et al., 2013) (C18:3, *P* < 8.85×10^-6^; Fig. 5b). Two caQTLs at chr9:140,450,305-140,450,725 and chr9:140,451,350-140,451,919 colocalized with eQTLs of *fad7* (Additional file 1: Fig. S4a). Peak SNPs (chr9.s_140450528 for chr9:140,450,305--140,450,725, chr9.s_140451649 for chr9:140,451,350--140,451,919) significantly influenced chromatin accessibility (Fig. 5c), the expression level of *fad7* (Fig. 5d) and the linolenic acid content (Fig. 5e). Additionally, the SNP chr9.s_140450528 within chr9:140,450,305-140,450,725 was predicted to disrupt the motifs of the MYB TF family (*P* < 5.86×10^-5^, Fig. 5f). Interestingly, the chromatin accessibility levels of the two caACRs were highly correlated (R^2^ = 0.67, *P* < 2.2×10^-16^; Additional file 1: Fig. S4b). Analysis of the ChIP-seq data of 104 TFs revealed that MYB, NAC, KNOX and COL TFs could simultaneously bind to both caACRs (Fig. 5g). These results suggest that this SNP affects MYB TF binding, alters chromatin accessibility and eventually leads to changes in fad7 expression, ultimately impacting the linolenic acid content.

We also identified a caACR located at chr2:10,495,708-10,496,333 that may regulate MADS TF 9 (*MADS9*). The peak SNPs (chr2.s_10496221 and chr2.s_10496222) of the caQTL of this caACR colocalized with the eQTL of *MADS*9 (Additional file 1: Fig. S5a, S5b). These two SNPs were predicted to disrupt binding motifs of the DOF TF family (*P* < 7.21×10^-5^, Additional file 1: Fig. S5c). *MADS9* has been identified as a candidate gene associated with flowering time (Xiao et al., 2021). Previous ChIP-seq has shown that this caACR contains a binding site for transcription factors involved in flowering, such as ZmELF3 and ZmCOL3 (Alter et al., 2016; Li et al., 2016) (Additional file 1: Fig. S5d). To validate the function of this gene, we edited the first exon of the gene and obtained two deletion mutants (Additional file 1: Fig. S5e). In the T1 generation mutants, we observed a trend toward delayed flowering compared with that of the wild type (WT) (Additional file 1: Fig. S5f).

## DISCUSSION

In this study, we utilized ATAC-seq technology to identify accessible chromatin regions in maize seedling leaves from a panel of 214 inbred lines and identified a total of 82,174 ACRs. Notably, more than 80% of these ACRs contain TF binding sites identified via ChIP-seq. Among these, 32,503 ACRs exhibited substantial variability in chromatin accessibility, ranging from 2-to 4-fold differences across the population. Since ATAC-seq is a tissue-specific assay and some functional SNPs could have been missed, we performed a genome-wide association study with ATAC-seq-only SNPs as well as imputed SNPs to explore the genetic basis of chromatin accessibility. The GWAS results revealed that imputation can increase statistical power, enabling the identification of more caACRs (10,835 vs 18,428, ATAC-seq-only SNPs vs imputed SNPs). There was no significant difference between the ATAC-seq-only and imputed SNPs in the proportion of peak SNPs disrupting TF motifs, indicating that imputation could be performed for chromatin accessibility GWASs to recover those missing SNPs.

Our GWAS identified 27,004 caQTLs associated with 18,428 caACRs via imputed SNPs. These caACRs demonstrated significantly greater average heritability than noncaACRs did. One plausible mechanism by which caQTLs may influence chromatin accessibility is through the disruption of TF binding sites. Our motifbreakR analysis predicted that over 44% of the peak SNPs within caACRs could disrupt TF motifs, particularly in the ERF, LBD, and TCP TF families. While previous studies in humans have shown that genetic variations in TF motifs can alter chromatin structure and establish chromatin accessibility (e.g., HNF, FOXA, and CEBP families; Mayran and Drouin 2018; Nagaki and Moriwaki 2008), similar studies in plants are lacking. In this study, our data showed that SNPs that potentially disrupted TF binding in plants could similarly influence chromatin accessibility.

We also integrated ATAC-seq data with other omics data to study how ACR could influence gene expression. A total of 1,990 pairs of caACRs and genes were identified, while 46.8% of the distal caACRs were more than 40 kb away from their target genes. These findings suggest that there are many long-range regulatory element interactions in the maize genome, which is consistent with previous research (Ricci et al., 2019). For example, we showed that the expression of *gpc4* could be regulated by proximal (<1 kb) and distal (117 kb) caACRs. We also identified two caACRs located at the 3’ end of fatty acid desaturase (*fad7*), which could influence chromatin accessibility by disrupting the TF binding motif, thereby affecting *fad7* expression and modifying the linolenic acid content. We also identified a caACR that can influence chromatin accessibility and the expression of *MADS9 simultaneously*, ultimately influencing flowering time. Compared with previous studies on the regulation of genes and traits (Li et al., 2013), our research revealed the potential genetic basis of altered chromatin accessibility in the maize population and shed light on the molecular mechanisms underlying the formation of complex traits.

The main limitation of this study is that ACRs are highly dynamic during cell differentiation and tissue development; however, our analysis explored only their regulatory potential in leaf tissue. Single-cell ATAC-seq is a promising solution to overcome this limitation (Pott and Lieb, 2015). By applying these technologies across multiple tissues, it is possible to systematically map the dynamic, cell type-specific changes in ACRs. Single-cell resolution cis-regulatory maps can be generated by combining single-cell ATAC-seq and RNA-seq data from six maize tissues to explore the relationships between CREs and TFs across different cell types and developmental stages (Marand et al., 2021). In Arabidopsis and rice root tissues, single-cell RNA and ATAC-seq revealed how chromatin accessibility influences gene expression and differentiation pathways (Farmer et al., 2021; Zhang et al., 2021). Recently, scATAC-seq revealed extensive chromatin accessibility variation linked to over 4.6 million genetic variants, primarily exhibiting cell type-specific effects (Marand et al., 2024). Taken together, population-scale chromatin accessibility analysis is a powerful tool for dissecting the regulatory potential of genetic variants associated with traits of interest. In the future, the application of single-cell technologies at the population level will be crucial for identifying key regulatory factors that determine cell fate and tissue development processes.

## MATERIALS AND METHODS

### Analysis of ATAC-seq data

The ATAC-seq data were analysed via a previously described method (Yan et al 2020). The analysis guidelines consisted of quality control, read mapping, and peak calling. Specifically, potential adapter sequences of the 214 ATAC-seq raw data were removed from the sequencing reads via fastp (Chen et al., 2018), and the quality of the sequencing data was then evaluated via FastQC (http://www.bioinformatics.babraham.ac.uk/projects/fastqc/). The cleaned reads were then aligned to the B73 V4 (Jiao et al., 2017) version of the reference genome via BWA v0.7.17 (Li 2013). The alignment reads were further filtered via SAMtools v1.13 (Danecek et al., 2021) to remove reads with low mapping quality (MAPQ < 30), which were mapped to chloroplasts and mitochondria, as well as PCR duplicates. Peak calling was performed via MACS2 v2.1.0 (Zhang et al., 2008). Duplicated reads were not considered (--keep-dup=1) during peak calling to achieve better specificity. The shifting size (--shift) used in the model was determined by the analysis of cross-correlation scores via the phantompeakqualtools (Landt et al 2012) package (https://code.google.com/p/phantompeakqualtools). The parameter “--call-summits” was used to call narrow peaks.

### Analysis of ChIP-seq data

Several datasets used in this study have already been published, including ChIP-seq data for 104 transcription factors (Tu et al., 2020) and histone modifications (H3K27ac, H3K27me3, H3K4me1, H3K4me3) (Peng et al., 2019). Using the same procedure as the aforementioned ATAC-seq analysis guidelines, quality control, alignment, and removal of reads with low mapping quality from the raw data were performed. Unlike ATAC-seq analysis, ATAC-seq identified narrow peaks, whereas ChIP-seq requires adjusting the parameters of MACS2 v2.1.0 to “--broad” for identifying broad peaks.

### Analysis of ChIA-PET data

**The** ChIA-PET data for RNA polymerase II and H3K4me3 used in this study were obtained from previously published research. The ChIA-PET data analysis pipeline utilized the ChIA-PET tools (Li et al., 2010) software package, including linker filtering, mapping tags to reference genomes, and identifying protein binding sites and chromatin interactions. To obtain more robust chromatin interactions, detected interactions were included only when the paired-end tag count was ≥ 3 and at least one interaction anchor overlapped with a peak in the corresponding ChIP-seq data.

### Quantification of ACRs and correction of batch effects

The ACRs identified from the 214 samples were merged via the default parameters of Bedtools v2.30.0 (Quinlan and Hall 2010). A total of 170,356 ACRs were obtained. The exclusion of ACRs with a frequency below 15 resulted in the retention of 82,174 ACRs in the final analysis. In ATAC-seq, the cut sites generated by the Tn5 enzyme are considered the most open regions. The cut-off site can be defined as the + 4 position for reads on the forward strand and the −5 position for reads on the reverse strand, taking into account the offset characteristic of the Tn5 enzyme. The number of cut sites in each ACR was counted across different samples and then subjected to median normalization by dividing it by the length of the ACR. Finally, the cut-off site count for each sample was adjusted to a value of 1 million to quantify the ACRs.

Since the ATAC-seq experiments were conducted in two batches, PCA was performed on the quantification matrix of the ACRs to investigate whether there was a noticeable batch effect. The results clearly revealed that the 214 samples were separated into two batches, indicating a significant batch effect. To address this, the quantification matrix of the ACRs was corrected via the “COMBAT” function from the R package SVA (Johnson et al 2007), with default parameters. After the correction, a PCA was conducted again, and the samples from the two batches were mixed together, indicating a reduction in the batch effect.

### Genotype identification and imputation

Using the BAM files generated from ATAC-seq data alignment, the standard pipeline of Sentieon v202112.06 (Kendig et al., 2019) software was used to genotype the samples. The pipeline consists of four main steps: removing or marking duplicates, indel realignment, base quality score recalibration, and variant calling. After genotyping each sample, we applied filtering via VCFtools v0.1.16 (Danecek et al 2011) (--minQ 30, --minGQ 10, --minDP 3) and merged the filtered SNPs via BCFtools v1.10.2 (Danecek et al 2021) merge. The merged SNP matrix served as the reference SNP set. Then, the Sentieon variant calling functionality was reapplied, and another round of merging was performed, resulting in a total of 349,492 SNPs.

To impute the genotypes, a reference set of 10 million high-quality SNPs from 507 published maize inbred lines was utilized. Beagle v0.27 (Browning et al., 2018) software was employed for genotype imputation. To assess the accuracy of imputation, we randomly selected 1% of the SNPs from the 340,000 SNPs identified in the 214 samples as evaluation sites. The remaining SNPs were imputed, and the genotypes of the 214 samples at the same SNP positions were extracted as the randomly selected evaluation sites. The imputation accuracy for each sample ranged from 91% to 98%, with an average of 96%. After imputation, a total of 10,770,214 SNPs were obtained. Further filtering was performed by retaining SNPs with a minor allele frequency (MAF) > 0.05 and a missing rate of less than 10%, resulting in approximately 7,443,172 SNPs.

### Calculation of heritability

To assess whether ACRs are genetically controlled, LDAK5 (Speed et al., 2012) software was used to estimate the heritability of 58,541 ACRs. First, the “ldak5 --cut-weights” and “--calc-weights-all” functions were used to calculate the weights of the SNPs. Then, on the basis of these weights, the “ldak5--calc-kins” function was used to calculate the kinship matrix. Finally, the heritability of each ACR was computed via the “ldak5 --reml” function.

### Genetic association analysis of ACRs and gene expression

A genome-wide association analysis was performed via the fastGWA-mlm function of GCTA v1.94.1 (Jiang et al., 2019). The set of 7,443,172 SNPs was used with a minor allele frequency (MAF) > 0.05 and a missing rate of less than 10% from the 21 materials. This analysis was conducted on 58,541 ACRs that were present in at least 90% of the materials. In the analysis, both the kinship matrix and the calculated population structure were incorporated via GCTA to account for relatedness and population stratification. The significant SNPs were determined by controlling the false discovery rate (FDR) at 0.05 via the Benjamini_Hochberg (BH) method (BH rejection threshold: P < 6.72×10^--9^).

Additionally, gene expression data from 197 maize inbred lines at the seedling stage and genotype data from 507 maize inbred lines from previously published studies were utilized. For the analysis of the 22,098 genes expressed at the seedling stage and the 10 million SNPs, a genome-wide association analysis was performed via fastGWA-mlm, with a significance threshold set at 1×10^-8^.

### Disruption of the Motif

A total of 5,571 SNPs were selected that showed the most significant associations with ACRs in the GWAS analysis, either by being located within the ACR or in strong linkage disequilibrium (LD > 0.7) with SNPs within the ACR. These SNPs were then subjected to motif disruption prediction via the R package motifBreakR (Coetzee et al., 2015). The significance threshold for motif disruption was set at 1×10^-4^. The motifBreakR analysis utilized the Jaspar2022 database (http://jaspar.genereg.net), from which 41 maize-specific TF motifs and 568 Arabidopsis motifs were filtered as references.

### Colocalization and association analysis of ACRs with gene expression

Two approaches were employed to obtain target genes for caACRs. The first approach involved colocalization analysis between peak SNPs in caQTLs and peak SNPs in eQTLs. Specifically, PLINK v1.9 (Purcell et al., 2007) was used to determine whether both peak SNPs were in strong linkage disequilibrium (LD). If *r^2^* > 0.7, it was considered evidence of colocalization between the ACR and the gene.

The second approach utilized association analysis between ACRs and gene expression to identify genes that might be regulated by ACRs. This analysis resembles a transcriptome-wide association study (TWAS). Since there was limited overlap (56 individuals) between the 214 individuals with ACR data and the 197 individuals with gene expression data, FUSION (Gusev et al., 2016) imputed the cis-regulatory effects of ACRs from the SNP genotype to the gene expression data of individuals without ACR data. Finally, association analysis between ACRs and gene expression was conducted via a significance threshold of 1×10^-6^.

### Enrichment analysis of traits

QTLs associated with agronomic traits and metabolic traits were collected from previously published methods (Gui et al., 2020). Agronomic traits were reclassified into flowering (days to anthesis, days to silking, days to tasselling, the tassel anthesis interval, the tassel silk interval, and the anthesis silk interval); morphology (leaf angle, leaf length, leaf width, plant height, main tassel branch length, branch number, ear leaf length, and the tassel silk interval); yield (100-grain weight, ear row number, kernel width, cob weight, kernel length, ear height, and ear weight); and metabolic traits were classified into carbohydrates (erythritol, fructose, fucose, galactinol, glucose, glycerol, raffinose, rhamnose, sucrose, trehalose, and xylose); and lipids (Oil-C16_0, Oil-C16_1, Oil-C18_0, Oil-C18_1, Oil-C18_2, Oil-C18_3, Oil-C20_0, Oil-C20_1, Oil-C22_0, Oil-C24_0, Oil-C160_C161, Oil-C160_C180, Oil-C180_C181, and Oil-C180_C200. Oil-C181_C182, Oil-C182_C183, Oil-C200_C201, Oil-C200_C220, Oil-C220_C240), amino acids (alanine, alanine beta, arginine, asparagine, aspartic acid. glutamine, glycine, histidine, homoserine, isoleucine, leucine, lysine, methionine, ornithine, phenylalanine, proline, serine, threonine, tryptophan, tyramine, tyrosine, valine). The number of QTLs for each category of traits will be counted, and then overlaps will be performed with the set of 1,555 ACRs. Additionally, 100 random samples of the same number and length as the 1,555 ACRs were extracted from the whole genome as controls. The overlaps with each category of trait QTLs were calculated, and the p value was calculated as the number of times the control overlaps exceeded the overlapping ACRs divided by 100.

### Data availability statement

The ATAC-seq data of 214 maize inbred lines is currently being uploaded to the NCBI database.

## ACKNOWLEDGEMENTS

We thank the high-throughput computing platform of the National Key Laboratory of Crop Genetic Improvement at Huazhong Agricultural University and H. Liu for computing support.

## AUTHOR CONTRIBUTIONS

Y.Z., S.Z., and N.Y. conceived the project. H.N., X.T. and K.G. generated ATAC-seq data. L.N. generated the *MADS9* mutants. Computational analyses were performed by Y.Z. and T.Z. and D.C. The manuscript was drafted by Y.Z. and reviewed and edited by L.Z., W.L., Y.X., J.Y., S.Z., and N.Y.

