## Additional_Figure_and_Table for "Pan-cistrome analysis of the leaf accessible chromatin regions of 214 maize inbred lines"

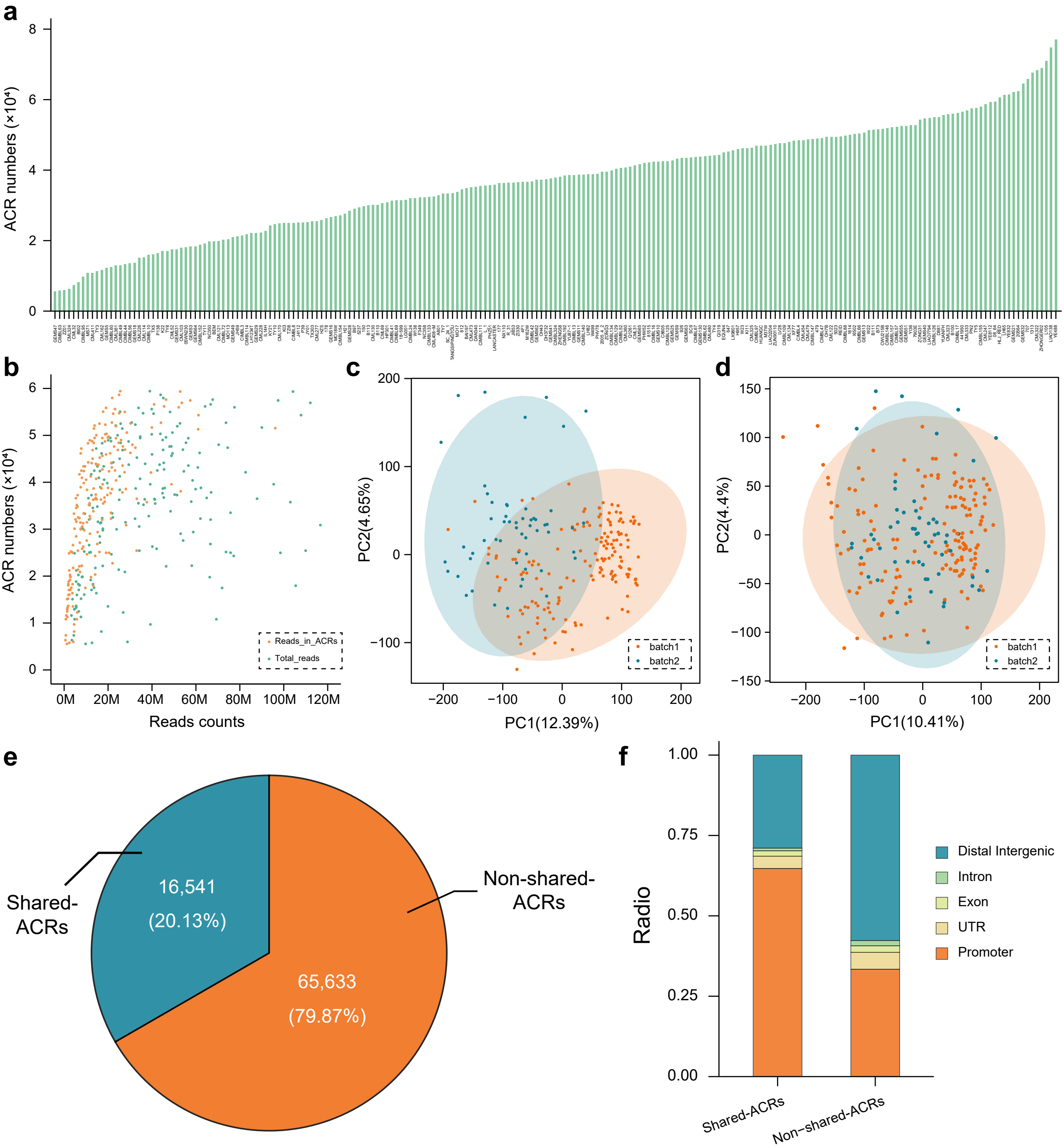
a
ACR numbers (×10⁴)
8
6
4
2
0
b
ACR numbers (×10⁴)
Reads counts
0M
20M
40M
60M
80M
100M
120M
Reads_in_ACRs
Total_reads
c
PC2(4.65%)
PC1(12.39%)
batch1
batch2
d
PC2(4.4%)
PC1(10.41%)
batch1
batch2
e
Shared-ACRs
16,541 (20.13%)
Non-shared-ACRs
65,633 (79.87%)
f
Radio
1.00
0.75
0.50
0.25
0.00
Distal Intergenic
Intron
Exon
UTR
Promoter
Shared-ACRs
Non−shared-ACRs

**Fig.S1** **a** The number of ACRs identified by 214 samples. **b** Scatterplot between the number of ACRs and reads counts, including cleaned reads and reads in ACRs. **c** PCA of 214 samples ACRs before correction. **d** PCA of 214 samples ACRs after correction. **e** Number and radio of shared-ACRs and non-shared-ACRs. **f** Bar graph showing the percentage of ACRs distribution among functional genomic elements.

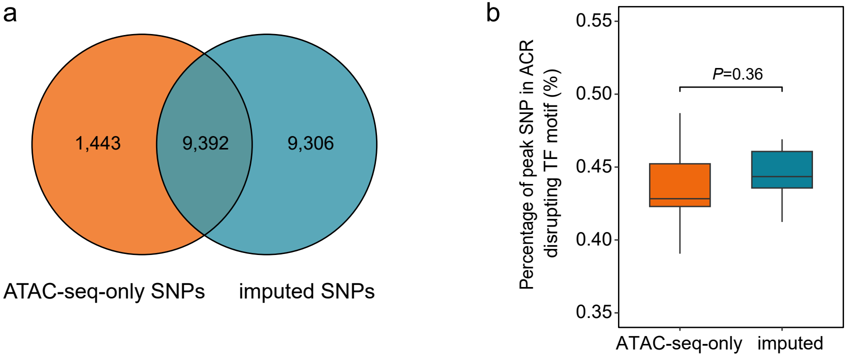
a
1,443
9,392
9,306
ATAC-seq-only SNPs
imputed SNPs
b
Percentage of peak SNP in ACR disrupting TF motif (%)
P=0.36
0.55
0.50
0.45
0.40
0.35
ATAC-seq-only
imputed

**Fig.S2 a** The number of ACRs with significant SNPs identified from ATAC-seq-only and imputed SNPs. **b** The proportion of peak SNPs within ACRs disrupting motifs identified between ATAC-seq-only and imputed SNPs.

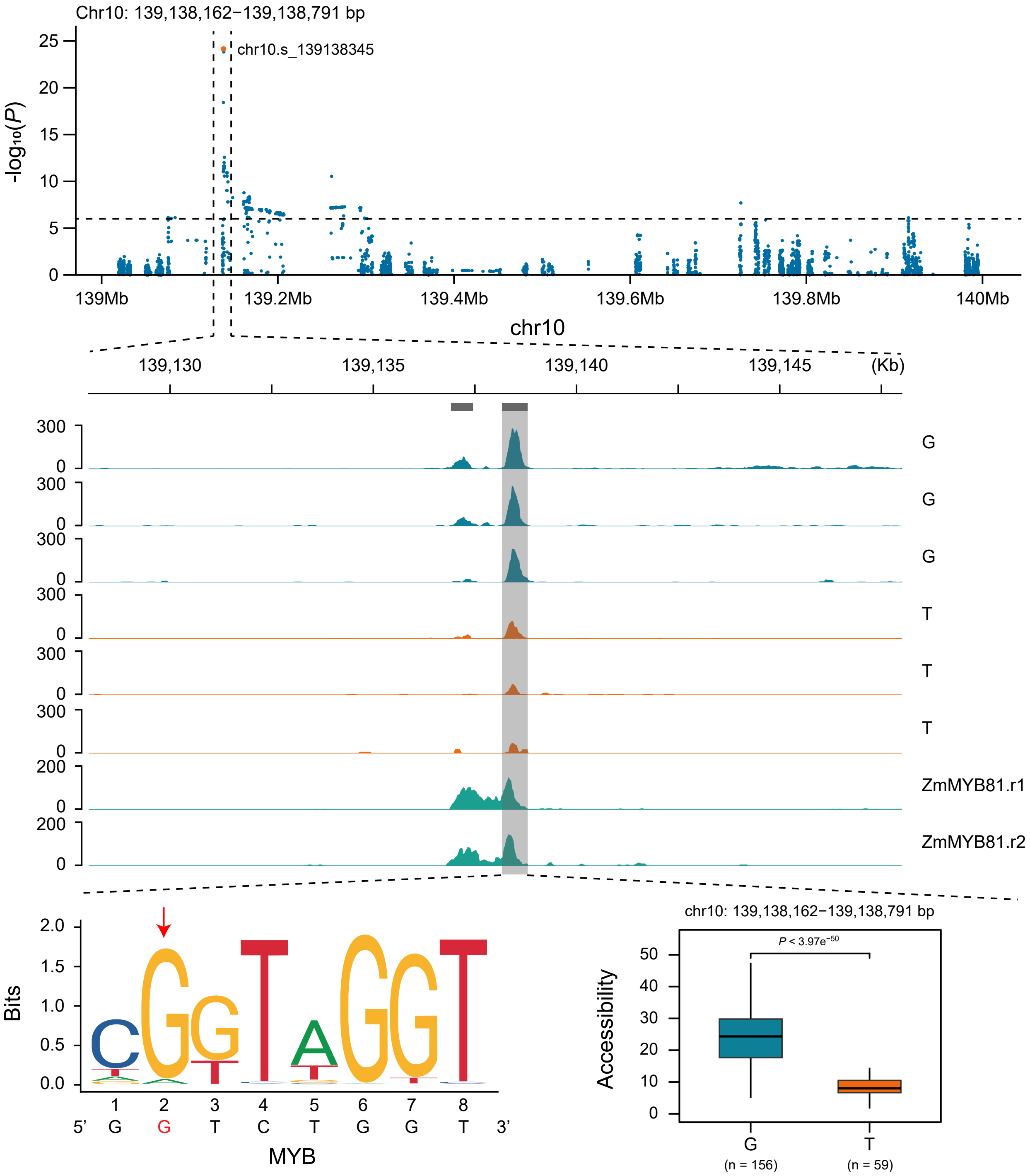

Chr10: 139,138,162−139,138,791 bp
-log10(P)
chr10.s_139138345
139Mb
139.2Mb
139.4Mb
139.6Mb
139.8Mb
140Mb
chr10
139,130
139,135
139,140
139,145
(Kb)
G
G
G
T
T
T
ZmMYB81.r1
ZmMYB81.r2
Bits
5' G G T C T G G T 3'
MYB
chr10: 139,138,162−139,138,791 bp
P < 3.97e−50
Accessibility
G
(n = 156)
T
(n = 59)

**Fig.S3** In the Manhattan plot, chr10.s_139136345 is the peak SNP within the region chr10:139,138,162-139,138,791 (top). The peak SNP is predicted to disrupt the MYB family motif with the red arrow (bottom left). The box plot displays the significant impact of the two genotypes on chromatin accessibility (bottom right). The genome browser track shows the difference of chromatin accessibility of two genotypes and the ChIP-seq data indicating ZmMYB81 binds to the ACR region (center).

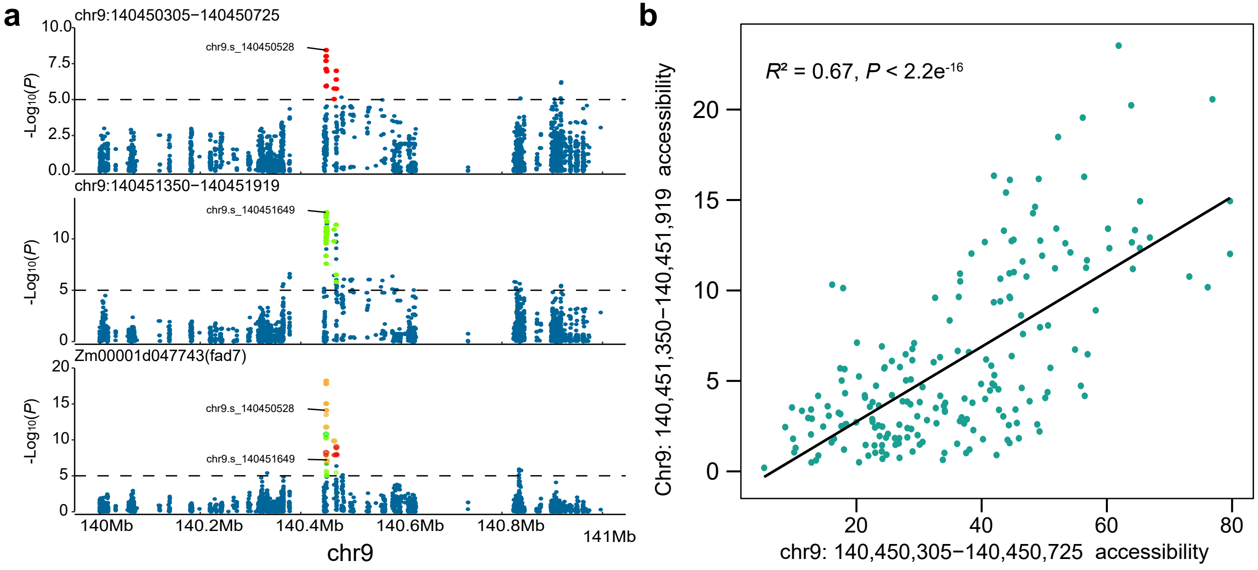

a
chr9:140450305−140450725
chr9.s_140450528
chr9:140451350−140451919
chr9.s_140451649
Zm00001d047743(fad7)
chr9.s_140450528
chr9.s_140451649
-Log10(P)
140Mb
140.2Mb
140.4Mb
140.6Mb
140.8Mb
141Mb
chr9
b
R² = 0.67, P < 2.2e-16
Chr9: 140,451,350−140,451,919 accessibility
chr9: 140,450,305−140,450,725 accessibility

**Fig.S4 a** Local Manhattan plots for chr9:140,450,305-140,450,725, chr9:140,451,350-140,451,919, and *fad7*. Red dots represent significant SNPs within chr9:140,450,305-140,450,725, green dots within chr9:140,451,350-140,451,919. Labels represent peak SNPs of ACRs. **b** The chromatin accessibility of chr9:140,450,305-140,450,725 and chr9:140,451,350-140,451,919 shows a significant correlation.

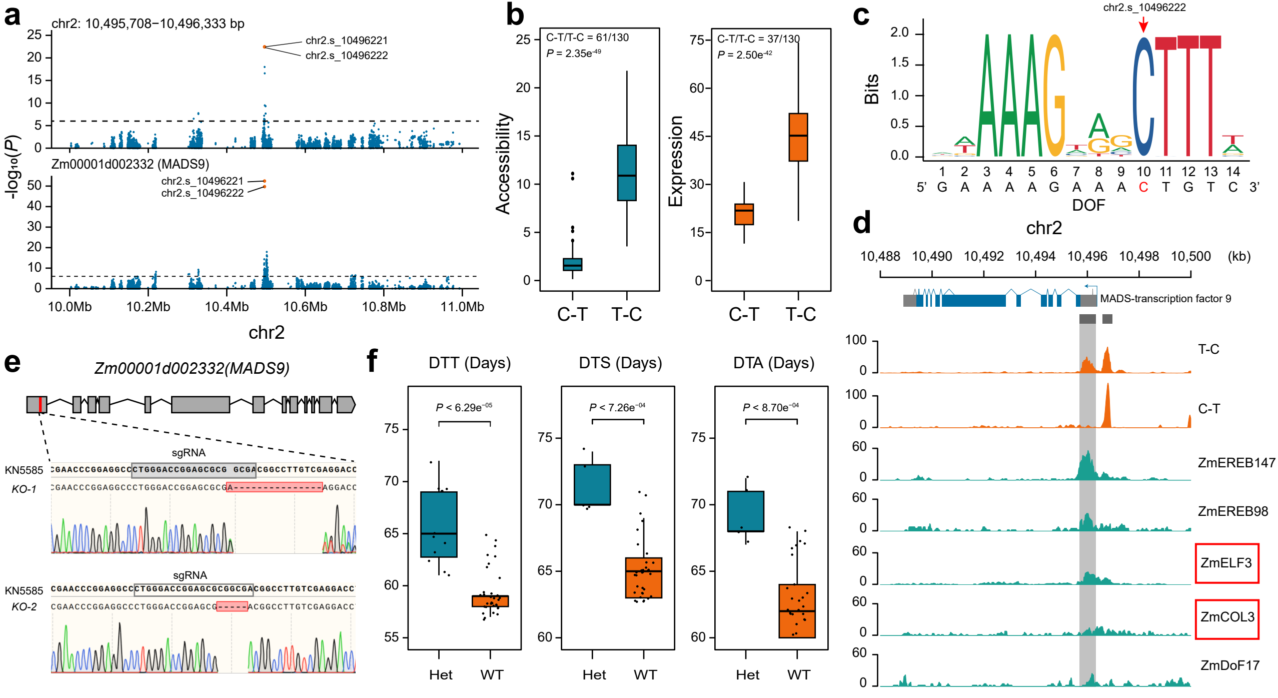

a
chr2: 10,495,708−10,496,333 bp
chr2.s_10496221
chr2.s_10496222
Zm00001d002332 (MADS9)
-log₁₀(P)
10.0Mb 10.2Mb 10.4Mb 10.6Mb 10.8Mb 11.0Mb
chr2
b
C-T/T-C = 61/130
P = 2.35e-49
Accessibility
C-T/T-C = 37/130
P = 2.50e-42
Expression
C-T T-C
c
chr2.s_10496222
Bits
5' G A A A A G A A A C T G T C 3'
DOF
d
chr2
10,488 10,490 10,492 10,494 10,496 10,498 10,500 (kb)
MADS-transcription factor 9
T-C
C-T
ZmEREB147
ZmEREB98
ZmELF3
ZmCOL3
ZmDoF17
e
Zm00001d002332(MADS9)
sgRNA
KN5585
KO-1
KO-2
f
DTT (Days)
DTS (Days)
DTA (Days)
P < 6.29e-05
P < 7.26e-04
P < 8.70e-04
Het WT

**Fig.S5 a** Local Manhattan plot of chr2:10,495,708-10,496,333 and *mads9*. The red dots represent the two peak SNPs within both ACR and gene, namely chr2.s_10496221 and chr2.s_10496222, which are in complete linkage disequilibrium. **b** Two haplotypes of peak SNPs significantly impact on chromatin accessibility and *MADS9* expression. **c** The peak SNPs (chr2.s_10496222) are predicted to disrupt motifs of the DOF family with the red arrows. **d** The genome browser track displays the chromatin accessibility (orange) of the two different genotypes, as well as the TF (green) binding to the ACR. **e** *MADS9* structure and editing sites within coding sequences (CDs) of the *MADS9*. **f** *mads9* mutant shows a tendency for delayed flowering compared to the wild type. Flowering phenotypes include Days To Tasseling (DTT), Days To Silking (DTS), and Days To Anthesis (DTA).

**Table S1 Basic mapping data, FRiP, TSS enrichment, and ACRs number of 214 samples.**

| Sample | Mapping rate | Cleaned reads | Reads in ACRs | FRiP | TSS enrichment | ACRs number |
|---|---|---|---|---|---|---|
| 150 | 93.54% | 17632884 | 9115846 | 0.51698 | 14.5 | 29768 |
| 177 | 89.37% | 25048828 | 13196289 | 0.52682 | 9.13 | 36266 |
| 178 | 90.75% | 40174586 | 8733778 | 0.2174 | 6.56 | 26291 |
| 479 | 85.34% | 20065822 | 11671980 | 0.58168 | 12 | 48952 |
| 647 | 83.18% | 33111838 | 16211527 | 0.4896 | 13.81 | 45254 |
| 707 | 83.53% | 61314320 | 43538479 | 0.71009 | 13.33 | 65858 |
| 812 | 91.25% | 13082312 | 5495576 | 0.42008 | 17.03 | 34526 |
| 926 | 86.51% | 18029728 | 9521469 | 0.5281 | 12.44 | 43394 |
| 1313 | 74.60% | 58153575 | 34988315 | 0.60165 | 17.46 | 67601 |
| 1614 | 78.14% | 24344786 | 14668436 | 0.60253 | 12.29 | 50028 |
| 5023 | 80.57% | 21030737 | 13888171 | 0.66037 | 17.61 | 49447 |
| 5237 | 91.84% | 58779703 | 15247829 | 0.25941 | 10.36 | 29384 |
| 8902 | 92.83% | 16920572 | 4560598 | 0.26953 | 28.35 | 8184 |
| 20564 | 86.81% | 72360810 | 42218484 | 0.58344 | 13.99 | 62345 |
| 81515 | 88.18% | 82605876 | 27499039 | 0.33289 | 13.64 | 42126 |
| 441950 | 86.41% | 66705150 | 40975056 | 0.61427 | 12.77 | 56599 |
| 05WN230 | 90.30% | 7034860 | 1878275 | 0.267 | 16.1 | 18045 |
| 08_64 | 84.18% | 39014220 | 25508907 | 0.65384 | 12.69 | 59416 |
| 18-599 | 83.30% | 27962678 | 10386104 | 0.37143 | 24.65 | 31465 |
| 2005_4_2 | 82.95% | 39191788 | 17546683 | 0.44771 | 16.28 | 39567 |
| 4F1 | 81.28% | 11041150 | 5330894 | 0.48282 | 22.1 | 36610 |
| 7903E | 86.45% | 30757145 | 21734156 | 0.70664 | 13.71 | 52729 |
| A801 | 92.33% | 38517668 | 11685587 | 0.30338 | 8.58 | 32792 |
| B100 | 83.59% | 77565243 | 52189787 | 0.67285 | 19.17 | 55981 |
| B110 | 85.44% | 8508508 | 4715943 | 0.55426 | 19.35 | 30104 |
| B111 | 87.45% | 26463502 | 13566669 | 0.51266 | 11.07 | 51453 |
| B113 | 58.01% | 23858995 | 7515750 | 0.31501 | 11.54 | 29974 |
| B73 | 92.77% | 240457008 | 96121185 | 0.39974 | 10.72 | 51538 |
| B97 | 82.85% | 53054457 | 11734759 | 0.22118 | 6.82 | 29032 |
| BAI197 | 85.28% | 15093100 | 6070033 | 0.40217 | 11.01 | 34828 |
| BEM | 85.69% | 8106122 | 2869765 | 0.35402 | 20 | 19797 |
| C8605 | 81.54% | 13714620 | 7841882 | 0.57179 | 14.94 | 42477 |
| CIMBL10 | 86.39% | 5807278 | 1967306 | 0.33877 | 16.23 | 15990 |
| CIMBL109 | 71.66% | 48764845 | 26767620 | 0.54891 | 13.38 | 47637 |
| CIMBL111 | 71.92% | 41451037 | 17771889 | 0.42874 | 12.69 | 35476 |
| CIMBL114 | 61.60% | 14215282 | 4115492 | 0.28951 | 16.59 | 21733 |
| CIMBL125 | 75.19% | 42426422 | 23173370 | 0.5462 | 15.6 | 42515 |
| CIMBL126 | 85.60% | 48236169 | 27600700 | 0.5722 | 14.22 | 54988 |
| CIMBL129 | 72.81% | 58489666 | 23655136 | 0.40443 | 14.41 | 40406 |
| CIMBL13 | 86.73% | 127458961 | 28090213 | 0.22039 | 6.74 | 38567 |
| CIMBL133 | 76.04% | 17322922 | 6976033 | 0.40271 | 11.89 | 32357 |
| CIMBL134 | 80.04% | 19634998 | 8952026 | 0.45592 | 12.06 | 39826 |
| CIMBL140 | 77.27% | 32558905 | 16963503 | 0.52101 | 13.64 | 38678 |
| CIMBL146 | 80.12% | 51514222 | 23578771 | 0.45771 | 13.48 | 51942 |
| CIMBL147 | 76.52% | 21356582 | 12704107 | 0.59486 | 13.81 | 48757 |
| CIMBL152 | 87.95% | 5481678 | 2036342 | 0.37148 | 15.18 | 18831 |
| CIMBL154 | 91.50% | 16276616 | 7123601 | 0.43766 | 15.22 | 27120 |
| CIMBL155 | 82.11% | 45145404 | 21899234 | 0.48508 | 11.55 | 57958 |
| CIMBL157 | 57.25% | 23484860 | 13883457 | 0.59117 | 16.41 | 52090 |
| CIMBL16 | 87.95% | 54514178 | 20433429 | 0.37483 | 9.73 | 42328 |
| CIMBL17 | 76.40% | 64191017 | 45497652 | 0.70879 | 18.68 | 56169 |
| CIMBL192 | 75.44% | 25318982 | 10420248 | 0.41156 | 13.17 | 38389 |
| CIMBL24 | 91.17% | 66692244 | 17297249 | 0.25936 | 8.06 | 32034 |

| | | | | | | |
|---|---|---|---|---|---|---|
| CIMBL3 | 89.63% | 24859740 | 6240119 | 0.25101 | 7.3 | 21365 |
| CIMBL324 | 85.55% | 66929677 | 26520028 | 0.39624 | 13.73 | 37839 |
| CIMBL4 | 64.34% | 25740283 | 13469161 | 0.52327 | 12.89 | 48396 |
| CIMBL41 | 88.77% | 16451082 | 8652892 | 0.52598 | 11.87 | 31272 |
| CIMBL42 | 81.46% | 45362968 | 18348689 | 0.40449 | 12.9 | 36743 |
| CIMBL44 | 91.04% | 5524710 | 1530550 | 0.27704 | 16.68 | 13300 |
| CIMBL47 | 84.02% | 44849228 | 25891173 | 0.57729 | 15.07 | 49010 |
| CIMBL49 | 86.24% | 4756002 | 1366254 | 0.28727 | 18.02 | 12938 |
| CIMBL52 | 84.73% | 18214955 | 9983551 | 0.5481 | 13.99 | 40540 |
| CIMBL56 | 90.31% | 19945218 | 3987462 | 0.19992 | 18.16 | 13540 |
| CIMBL57 | 80.79% | 32424658 | 18771247 | 0.57892 | 15.34 | 52282 |
| CIMBL60 | 85.77% | 6655750 | 1397253 | 0.20993 | 11.72 | 12471 |
| CIMBL62 | 83.19% | 36869060 | 14460291 | 0.39221 | 8.88 | 43938 |
| CIMBL63 | 84.61% | 12681018 | 2398927 | 0.18917 | 18.29 | 5784 |
| CIMBL66 | 85.74% | 43814822 | 19304387 | 0.44059 | 12.13 | 50328 |
| CIMBL67 | 51.62% | 29405350 | 15564600 | 0.52931 | 15.14 | 43746 |
| CIMBL69 | 87.90% | 16831610 | 7379318 | 0.43842 | 13.18 | 31418 |
| CIMBL72 | 64.91% | 6492216 | 2664836 | 0.41047 | 19.67 | 20242 |
| CIMBL8 | 87.11% | 11520362 | 3882198 | 0.33699 | 13.28 | 25023 |
| CIMBL88 | 74.68% | 34646098 | 20275465 | 0.58522 | 17.15 | 49727 |
| CIMBL95 | 92.74% | 39282934 | 4540540 | 0.11559 | 9.18 | 9700 |
| CIMBL97 | 88.22% | 26591282 | 12958432 | 0.48732 | 11.13 | 46859 |
| CML103 | 89.87% | 105430954 | 13252706 | 0.1257 | 3.78 | 17951 |
| CML114 | 91.42% | 7699994 | 1783204 | 0.23159 | 10.58 | 15183 |
| CML121 | 83.54% | 8350438 | 2799808 | 0.33529 | 14.12 | 19866 |
| CML122 | 82.99% | 34613504 | 17397973 | 0.50264 | 13.23 | 49436 |
| CML130 | 88.17% | 17630862 | 7556665 | 0.4286 | 12.54 | 30101 |
| CML134 | 74.18% | 70382838 | 31088726 | 0.44171 | 13.1 | 47999 |
| CML162 | 76.54% | 3310018 | 965730 | 0.29176 | 20 | 11603 |
| CML170 | 66.98% | 4735972 | 2228616 | 0.47057 | 22.17 | 24859 |
| CML226 | 64.42% | 3800882 | 1347565 | 0.35454 | 25.36 | 15090 |
| CML228 | 91.63% | 58123301 | 8035712 | 0.13825 | 3.66 | 22217 |
| CML247 | 88.12% | 43935949 | 25036168 | 0.56983 | 16.96 | 58640 |
| CML277 | 88.70% | 109956802 | 33012233 | 0.30023 | 15.47 | 25429 |
| CML31 | 86.01% | 25299044 | 12094379 | 0.47806 | 14.95 | 41274 |
| CML32 | 73.86% | 4393830 | 786201 | 0.17893 | 15.62 | 7374 |
| CML322 | 87.37% | 116477432 | 63165532 | 0.5423 | 15.81 | 68334 |
| CML323 | 77.22% | 33876145 | 19045868 | 0.56222 | 13.03 | 55852 |
| CML324 | 76.92% | 3514238 | 583240 | 0.16596 | 10.77 | 6347 |
| CML325 | 70.56% | 61682179 | 30156962 | 0.48891 | 13.48 | 46260 |
| CML333 | 84.85% | 112102880 | 53575669 | 0.47792 | 16.54 | 56929 |
| CML360 | 86.26% | 34578610 | 14553782 | 0.42089 | 14.68 | 40734 |
| CML361 | 86.88% | 4614672 | 1440313 | 0.31212 | 17.31 | 12928 |
| CML408-M | 83.88% | 22164287 | 6339480 | 0.28602 | 8.55 | 32501 |
| CML411 | 67.70% | 4789976 | 783535 | 0.16358 | 10.06 | 10811 |
| CML454 | 88.31% | 55466114 | 24873831 | 0.44845 | 14.13 | 48421 |
| CML473 | 85.74% | 11534310 | 5772356 | 0.50045 | 12.94 | 35124 |
| CML479 | 82.97% | 29347942 | 18604558 | 0.63393 | 14.36 | 48697 |
| CML480 | 78.29% | 21614840 | 12524326 | 0.57943 | 13.16 | 44084 |
| CML52 | 86.31% | 71914047 | 9642937 | 0.13409 | 3.76 | 17478 |
| CML69 | 90.18% | 34166917 | 8196929 | 0.23991 | 6.43 | 30567 |
| CWU215B | 85.89% | 90024490 | 39943966 | 0.4437 | 9.78 | 51671 |
| D881 | 85.88% | 25478880 | 17742480 | 0.69636 | 18.53 | 55051 |
| DAN340 | 84.34% | 36104986 | 11272500 | 0.31221 | 10.55 | 35205 |
| DH3732 | 91.46% | 50308932 | 14413348 | 0.2865 | 10.39 | 37373 |
| EQUN4 | 85.77% | 25285156 | 12963862 | 0.51271 | 14.29 | 45022 |
| F349 | 49.97% | 32293806 | 11802553 | 0.36547 | 16.8 | 32239 |
| GEMS10 | 84.28% | 16295184 | 8965155 | 0.55017 | 13.05 | 42389 |

| | | | | | | |
|---|---|---|---|---|---|---|
| GEMS11 | 87.01% | 44958927 | 13387810 | 0.29778 | 9.31 | 38668 |
| GEMS13 | 73.10% | 49301302 | 28521596 | 0.57852 | 13.31 | 50588 |
| GEMS130 | 84.38% | 43458068 | 14689264 | 0.33801 | 10.08 | 43756 |
| GEMS16 | 90.36% | 11382498 | 4008524 | 0.35217 | 10.11 | 26603 |
| GEMS18 | 57.84% | 5465242 | 1882185 | 0.34439 | 22.75 | 13651 |
| GEMS21 | 82.63% | 93603844 | 55252974 | 0.59029 | 13.33 | 62200 |
| GEMS25 | 72.16% | 45495778 | 17419169 | 0.38287 | 12.01 | 42738 |
| GEMS28 | 85.17% | 16197058 | 5464780 | 0.33739 | 14.61 | 28460 |
| GEMS29 | 95.66% | 17568104 | 7970235 | 0.45368 | 24.23 | 22172 |
| GEMS3 | 83.39% | 13135710 | 3834681 | 0.29193 | 14.64 | 18275 |
| GEMS31 | 93.69% | 13637720 | 3255655 | 0.23872 | 11.39 | 17511 |
| GEMS32 | 78.73% | 55508263 | 35364328 | 0.6371 | 15.26 | 64470 |
| GEMS40 | 77.12% | 40714150 | 22307306 | 0.5479 | 12.58 | 54620 |
| GEMS41 | 56.75% | 44806307 | 21152101 | 0.47208 | 15.24 | 37670 |
| GEMS47 | 93.20% | 9765326 | 1262035 | 0.12924 | 13.2 | 5554 |
| GEMS49 | 89.54% | 27650452 | 7228849 | 0.26144 | 17.6 | 20883 |
| GEMS5 | 89.07% | 26887734 | 5955338 | 0.22149 | 13.1 | 12251 |
| GEMS51 | 92.41% | 52749736 | 30206909 | 0.57265 | 15.13 | 52553 |
| GEMS55 | 75.15% | 25737943 | 16086625 | 0.62502 | 14.26 | 52494 |
| GEMS62 | 90.28% | 12437804 | 6113313 | 0.49151 | 18.14 | 37246 |
| GEMS63 | 92.69% | 15141772 | 8828611 | 0.58306 | 20.78 | 43394 |
| GEMS64 | 80.36% | 6430602 | 1909851 | 0.29699 | 17.11 | 18306 |
| GEMS65 | 92.36% | 12340572 | 6957102 | 0.56376 | 12.13 | 41696 |
| GEMS9 | 93.84% | 60865913 | 21710809 | 0.3567 | 12.5 | 43155 |
| H21 | 91.95% | 54184583 | 10268710 | 0.18951 | 5.91 | 27575 |
| H957 | 83.42% | 40019970 | 19091619 | 0.47705 | 12.3 | 45941 |
| HLJ_RED | 83.64% | 131913712 | 85721740 | 0.64983 | 17.67 | 60556 |
| HP301 | 89.70% | 116659317 | 28212993 | 0.24184 | 6.85 | 30867 |
| HUANGC | 90.76% | 45082057 | 24445574 | 0.54225 | 27.06 | 46895 |
| HZS | 89.83% | 42117736 | 8167521 | 0.19392 | 5.12 | 25824 |
| IL14H | 88.37% | 138325162 | 22039342 | 0.15933 | 6.94 | 22743 |
| J4112 | 91.12% | 12100992 | 3575126 | 0.29544 | 7.86 | 25093 |
| JI853 | 86.61% | 41050468 | 14094081 | 0.34334 | 9.93 | 36438 |
| JY01 | 90.34% | 9145072 | 3027654 | 0.33107 | 15.95 | 25237 |
| K22 | 89.65% | 12464856 | 2834181 | 0.22737 | 9.86 | 17047 |
| KI11 | 87.70% | 49304247 | 20951651 | 0.42495 | 14.13 | 46163 |
| KI3 | 89.44% | 98462504 | 20481852 | 0.20802 | 12.18 | 24960 |
| KY21 | 88.52% | 77194011 | 14110748 | 0.1828 | 4.18 | 24302 |
| L_1 | 81.11% | 33816088 | 11606498 | 0.34322 | 12.84 | 35531 |
| L105 | 84.34% | 107716154 | 81332503 | 0.75506 | 15.03 | 70987 |
| LANCASTER | 87.83% | 57879001 | 18527705 | 0.32011 | 11.64 | 35874 |
| LG001 | 89.34% | 15385720 | 5590958 | 0.36339 | 15.79 | 31580 |
| LH1234T | 92.73% | 46447900 | 6887709 | 0.14829 | 2.92 | 22083 |
| LH65 | 87.08% | 64059449 | 44656741 | 0.69711 | 13.49 | 61348 |
| LH82 | 87.69% | 45774687 | 16879652 | 0.36876 | 10.92 | 38774 |
| LIAO138 | 86.50% | 99178798 | 72239449 | 0.72838 | 16.23 | 74769 |
| LIAO2204 | 80.36% | 14611966 | 7808380 | 0.53438 | 15.33 | 47191 |
| LIAO7794 | 86.44% | 35945921 | 25062888 | 0.69724 | 14.14 | 54712 |
| LV28 | 87.57% | 48010794 | 22749583 | 0.47384 | 12.95 | 47619 |
| LX9801 | 89.90% | 48876751 | 17416087 | 0.35633 | 9.59 | 45642 |
| M162W | 91.06% | 64907502 | 14350521 | 0.22109 | 9.76 | 36664 |
| M37W | 88.77% | 54076840 | 27713371 | 0.51248 | 18.98 | 46947 |
| MO113 | 93.64% | 128013784 | 14102419 | 0.11016 | 1.94 | 20335 |
| MO17 | 88.70% | 48472549 | 13295329 | 0.27429 | 10.39 | 33819 |
| MO18W | 89.11% | 66605457 | 15072939 | 0.2263 | 10.99 | 26836 |
| MS71 | 92.45% | 69296333 | 5766833 | 0.08322 | 1.62 | 10798 |
| NC350 | 89.66% | 66300306 | 12230100 | 0.18447 | 6.02 | 19770 |
| NC358 | 89.36% | 62061382 | 21357551 | 0.34414 | 15.06 | 32247 |

| | | | | | | |
|---|---|---|---|---|---|---|
| NX110 | 88.97% | 41313773 | 16763649 | 0.40576 | 15.24 | 36324 |
| OH43 | 52.79% | 51371084 | 19668894 | 0.38288 | 13.18 | 37282 |
| OH7B | 88.62% | 56540517 | 23581843 | 0.41708 | 14.06 | 49394 |
| P138 | 59.19% | 14644570 | 3745917 | 0.25579 | 16.92 | 16363 |
| P39 | 88.70% | 73668165 | 16051491 | 0.21789 | 10.18 | 25137 |
| PHV78 | 67.04% | 16478000 | 7541192 | 0.45765 | 12.29 | 38945 |
| PHW52 | 92.87% | 121734465 | 39536582 | 0.32478 | 7.5 | 42073 |
| PHZ51 | 88.46% | 103010340 | 37652235 | 0.36552 | 9.81 | 35766 |
| PN2 | 87.28% | 97432604 | 57413503 | 0.58926 | 14.95 | 57444 |
| Q1261 | 86.53% | 15804984 | 8105329 | 0.51283 | 13.66 | 40943 |
| QI319 | 86.43% | 22370274 | 10953279 | 0.48964 | 11.25 | 44239 |
| R_31 | 86.43% | 36692346 | 14839461 | 0.40443 | 16.69 | 36364 |
| REID | 90.85% | 46379110 | 19863648 | 0.42829 | 15.31 | 49577 |
| SC_24_1 | 75.69% | 21880792 | 8886882 | 0.40615 | 15.85 | 33362 |
| SC9 | 58.60% | 56575593 | 26612858 | 0.47039 | 13.41 | 43552 |
| ANGSIPINGTO | 89.26% | 75760423 | 21297650 | 0.28112 | 14.5 | 33502 |
| TX303 | 90.12% | 34244689 | 6262112 | 0.18286 | 7 | 25419 |
| TX5 | 66.70% | 9141874 | 2371344 | 0.25939 | 14.69 | 16054 |
| TY10 | 94.01% | 21525940 | 10393134 | 0.48282 | 22.77 | 24654 |
| TY11 | 88.45% | 12096968 | 2635947 | 0.2179 | 9.74 | 19170 |
| TY3 | 80.25% | 7642036 | 1483064 | 0.19407 | 12.33 | 11300 |
| TY4 | 87.89% | 45331609 | 21481163 | 0.47387 | 18.14 | 44141 |
| TY5 | 81.95% | 67641548 | 46369797 | 0.68552 | 15.54 | 57621 |
| TY7 | 84.79% | 9439042 | 4458636 | 0.47236 | 18.31 | 33309 |
| TY8 | 83.78% | 6667112 | 2151085 | 0.32264 | 16.32 | 17059 |
| TZI8 | 90.35% | 75792489 | 19145315 | 0.2526 | 12.02 | 24975 |
| W138 | 85.89% | 11378830 | 5155875 | 0.45311 | 19.69 | 32069 |
| W22 | 91.46% | 134534432 | 61010435 | 0.45349 | 19.29 | 51287 |
| W23 | 84.12% | 17041904 | 7567020 | 0.44402 | 12.51 | 46153 |
| W499 | 88.14% | 62280342 | 19990985 | 0.32098 | 10.57 | 38800 |
| XF77 | 85.04% | 17306980 | 10864988 | 0.62778 | 16.48 | 48314 |
| XI502 | 86.74% | 39896220 | 18704799 | 0.46884 | 14.18 | 50222 |
| YE488 | 83.51% | 148792194 | 113957920 | 0.76589 | 17.29 | 77030 |
| YE8112 | 91.07% | 104200390 | 52864460 | 0.50733 | 11.13 | 59333 |
| YE832 | 84.40% | 63236986 | 33710974 | 0.53309 | 10.22 | 61397 |
| YI36 | 87.07% | 73685338 | 44063448 | 0.59799 | 18.1 | 52686 |
| YU87-1 | 92.07% | 59878002 | 21788177 | 0.36388 | 11.56 | 38533 |
| YUANFH | 84.54% | 39878601 | 23172665 | 0.58108 | 16.77 | 55600 |
| ZHENG58 | 91.98% | 167489560 | 53093342 | 0.31699 | 12 | 38080 |
| ZHONGER02 | 87.43% | 106000670 | 63523473 | 0.59927 | 13.21 | 68985 |
| ZI330 | 84.09% | 87837919 | 33083580 | 0.37664 | 14.98 | 36496 |
| ZONG3 | 84.48% | 95412609 | 48876663 | 0.51227 | 21.96 | 39574 |
| ZONG31 | 88.03% | 107870520 | 61152528 | 0.56691 | 18.17 | 54305 |
| ZUN90110 | 84.81% | 74459948 | 32447842 | 0.43578 | 11.32 | 47370 |
| ZZ01 | 85.58% | 28835342 | 4162874 | 0.14437 | 17.71 | 5981 |

**Table S2 The distribution of ACRs in the population material.**

| Identity ACRs radio in sample | ACRs count |
|---|---|
| 100% (215) | 16541 |
| 90% (193) | 58541 |
| 80% (172) | 68238 |
| 50% (107) | 80350 |
| Total OCRs | 82174 |

**Table S3 The number of target genes for ACRs.**

| One ACR target gene | ACR counts | Ratio |
|---|---|---|
| 1 | 1227 | 78.91% |
| 2 | 248 | 15.95% |
| 3 | 66 | 4.24% |
| >4 | 14 | 0.90% |
